# Systematic identification of inter-chromosomal interaction networks supports the existence of RNA factories

**DOI:** 10.1101/2023.09.21.558852

**Authors:** Borislav Hrisimirov Hristov, William Stafford Noble, Alessandro Bertero

## Abstract

Most studies of genome organization have focused on intra-chromosomal (*cis*) contacts because they harbor key features such as DNA loops and topologically associating domains. Inter-chromosomal (*trans*) contacts have received much less attention, and tools for interrogating potential biologically relevant *trans* structures are lacking. Here, we develop a computational framework to identify sets of loci that jointly interact in *trans* from Hi-C data. This method, trans-C, initiates probabilistic random walks with restarts from a set of seed loci to traverse an input Hi-C contact network, thereby identifying sets of *trans*-contacting loci. We validate trans-C in three increasingly complex models of established *trans* contacts: the *Plasmodium falciparum var* genes, the mouse olfactory receptor “Greek islands”, and the human RBM20 cardiac splicing factory. We then apply trans-C to systematically test the hypothesis that genes co-regulated by the same *trans*-acting element (i.e., a transcription or splicing factor) co-localize in three dimensions to form “RNA factories” that maximize the efficiency and accuracy of RNA biogenesis. We find that many loci with multiple binding sites of the same transcription factor interact with one another in *trans*, especially those bound by transcription factors with intrinsically disordered domains. Similarly, clustered binding of a subset of RNA binding proteins correlates with *trans* interaction of the encoding loci. These findings support the existence of *trans* interacting chromatin domains (TIDs) driven by RNA biogenesis. Trans-C provides an efficient computational framework for studying these and other types of *trans* interactions, empowering studies of a poorly understood aspect of genome architecture.

## Introduction

Mammalian interphase chromosomes are exquisitely folded in three dimensions to enable precise regulation of gene expression (reviewed in (Hafner and Boettiger, 2023)). The study of such organization has been greatly advanced by sequencing-based chromosome conformation capture (3C) technologies, chiefly Hi-C (Lieberman-Aiden et al., 2009), and by orthogonal imaging approaches. A rapidly growing body of evidence indicates that while a large portion of 3D genome architecture is largely invariant across cell types, specific dynamic changes play a critical role in regulating gene expression during development and in disease (Krumm and Duan, 2019; Zheng and Xie, 2019).

Most of our current understanding of 3D genome architecture centers around chromatin folding within individual chromosomes, that is, on intra-chromosomal or *cis* contacts. These contacts give rise to a variety of hierarchical features at different genomic scales, including different types of DNA loops (i.e., cohesin-mediated looping and promoter-enhancer pairing), topologically associating domains (TADs; sub-megabase domains of preferential self-interaction) (Dixon et al., 2012), and A/B compartments (chromosome-wide segregation of active/inactive chromatin resulting from inter-TAD interactions) (Lieberman-Aiden et al., 2009). In contrast, interactions across different chromosomes (inter-chromosomal or *trans* contacts) are poorly understood.

Chromosome-wide *trans* genome architecture distinguishes non holocentric chromosomes of eukaryotic species in either a type-I, Rabl-like configuration (i.e., featuring centromere clustering, telomere clustering, and/or a telomere-to-centromere axis) or a type-II configuration characterized by chromosome territories (Hoencamp et al., 2021). The latter is typical of mammalian chromosomes, which tend to occupy distinct domains of the interphase nucleus (Cremer and Cremer, 2010). Although chromosome territories limit the possibility for *trans* contacts, they do not represent hard boundaries: regions that overcome territorial topological restrictions engage with each other within mRNA and tRNA factories, polycomb domains, the nucleolus, nuclear speckles, and potentially other nuclear subcompartments (Fig. 1A; (Bhat et al., 2021)). Some of these *trans* contacts involve specific loci whose interactions are important to gene regulation in enhancer hubs (Monahan et al., 2019), transcription factories (Osborne et al., 2004, 2007; Papantonis et al., 2012), and splicing factories (Bertero et al., 2019). Despite these and a few other examples, whose discovery was serendipitous or informed by domain-specific prior knowledge, the systematic discovery of functional *trans* interactions is currently very challenging.

**Figure 1:**
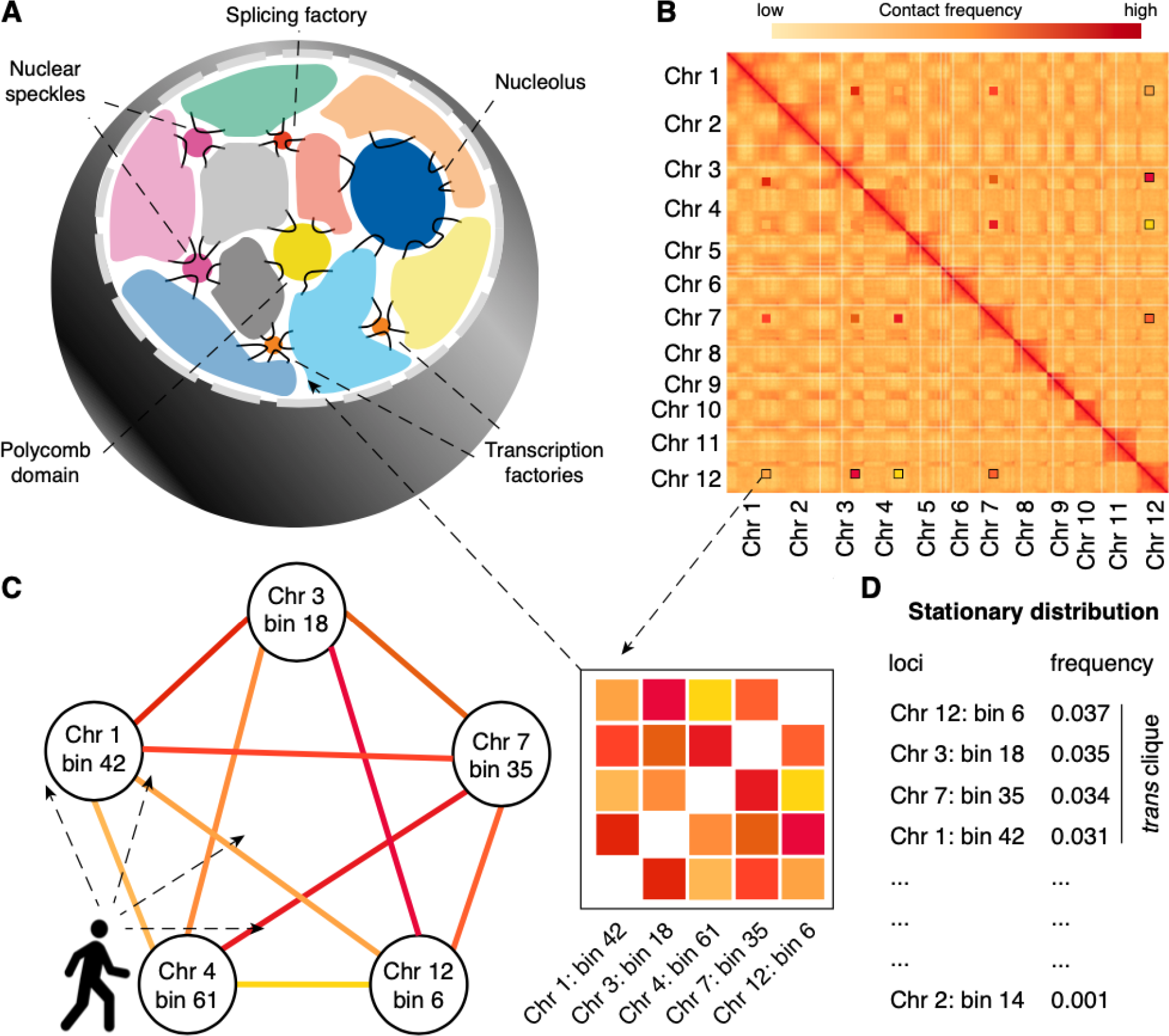
**(A)** Schematic of typical inter-chromosomal genome organization in mammals. Inter-chromosomal interactions mainly involve genomic domains that extrude from chromosome territories and engage with a variety of membraneless structures involved in gene regulation. **(B)** A Hi-C matrix captures the contact frequency of loci in a genome-wide fashion. Zoomed in boxes illustrate that specific loci located on different chromosomes can exhibit strong *trans* contacts among themselves. **(C)** Trans-C employs a random walk algorithm that traverses the Hi-C contact graph choosing to move to a node (bin) probabilistically based on the strength of the edge (interaction). **(D)** The output is a list of loci ranked by how frequently they are visited during the random walk: more frequently visited loci interact more strongly in *trans*.

One reason for this difficulty is that *trans* contacts are 5 to 10 times less frequent than *cis* contacts, depending on cell type and assay type. The number of possible *trans* contacts is also much larger than the number of possible *cis* contacts; thus, *trans* contact data is typically quite sparse. Most importantly, there is a lack of robust statistical and computational approaches to confidently identify reproducible *trans* contacts. In this manuscript, we overcome this limitation by providing a computational framework that systematically finds sets of jointly interacting loci from Hi-C data.

The method, trans-C, takes as input a Hi-C contact map as well as, optionally, one or more more seed loci and uses a random walk algorithm to identify sets of *trans*-contacting loci. We validate trans-C in three established problems of increasing complexity in *Plasmodium*, mouse, and human. We then deploy trans-C to systematically test the hypothesis that genes co-regulated by the same *trans*-acting element (i.e., a transcription or splicing factor) co-localize in 3D to form “RNA factories” that maximize the efficiency and accuracy of RNA biogenesis (Bertero, 2021). Our results suggest that many transcription factors (TFs) participate in *trans* interacting chromatin domains (TIDs), and that these types of interactions are particularly common for TFs with intrinsically disordered domains; in contrast, only a subset of RNA binding proteins (RBPs) are enriched at TIDs, but these enrichments can be exceptionally strong and potentially disease-relevant.

Overall, trans-C provides a powerful way to uncover and measure various types of *trans* interactions, empowering both discovery and hypothesis-driven studies of genome structure-function relationships.

## Results

### trans-C randomly walks the Hi-C graph

Our goal is to algorithmically identify, from a given set of Hi-C data, a collection of genomic loci that exhibit strong *trans* interactions. We represent the Hi-C data as a matrix, referred to as a “contact map”, in which each axis corresponds to the complete genome and entries in the matrix represent Hi-C contact counts (Fig. 1B). In practice, the genomic axes are discretized using fixed-width bins. The bin size is thus inversely proportional to the effective resolution of the contact map. The contact map can be thought of as the adjacency matrix of a corresponding Hi-C graph, in which nodes are genomic loci (bins) and edges are weighted by the corresponding contact counts (Fig. 1C). Our goal is thus to find dense subgraphs in this Hi-C graph.

The problem of dense subgraph discovery arises in many application domains and consequently has been very widely studied. Depending on the exact formulation and the notion of density, theoretical computer science has shown that the problem complexity ranges from easily solved in linear time via a max flow algorithm (Khuller and Saha, 2009) to computationally intractable (NP-hard) (Charikar, 2000). Common techniques to approximate the latter case, to which our specific problem belongs, are greedy approaches, which iteratively select the best option available at the moment without guaranteeing that this strategy will bring the global optimal result, (Charikar, 2000) and semi-random models, which account for model errors by incorporating both adversarial and random choices in instance generation (Bhaskara et al., 2010).

trans-C approaches the discovery task of finding strongly interacting loci in *trans* using a random walk with restart algorithm (Fig. 1C). This general approach has been applied successfully in domains as diverse as web searching (Gibson et al., 2005), protein remote homology detection (Weston et al., 2004), and gene functional prediction (Mostafavi et al., 2008). Prior to the random walk operation, trans-C performs three pre-processing steps on the provided Hi-C contact map. First, to control for sequencing and accessibility biases, Hi-C counts are ICE-normalized (Imakaev et al., 2012), thus equalizing the sum of counts per row and column in the matrix. Second, the resulting matrix is processed using a binomial model to estimate interaction p-values based on an empirical null model that accounts for biases arising from chromosomal territorialization (i.e., small, gene-rich chromosomes generally occupying the nuclear interior and interacting more with each other than with large, gene-poor chromosomes). Third, the negative log p-values are used as weights for the network edges and subsequently refined using a “donut filter” (Rao et al., 2014) to highlight points that are local maxima. trans-C then carries out a random walk with restart algorithm on the pre-processed Hi-C graph. Each walk is initiated from a randomly selected locus and moves from a node to a neighboring one probabilistically based on the weight of the edge. A parameter *α* controls the probability that the walk will restart at a new, randomly selected locus. Mathematically, as an infinite number of walks are performed, the frequency with which each node is visited converges to a stationary distribution. This distribution can be computed analytically using the Perron-Frobenius theorem. We use the stationary distribution to obtain a ranked list of *trans* interacting bins (Fig. 1D), because the most frequently visited nodes are the ones that interact most strongly with the seed loci.

### Trans-C uncovers the clustering of *var* genes critical for *P. falciparum* immune evasion

Having developed trans-C, we set out to test its ability to uncover known sets of loci that interact together in *trans* in three different organisms. First, we focused on the protozoan *Plasmodium falciparum*, the parasite responsible for the most lethal form of malaria, which has a haploid nuclear genome of 22.9 Mb across 14 chromosomes.

The three-dimensional organization of the *P. falciparum* genome is strongly associated with gene expression (Ay et al., 2014), particularly for genes involved in pathogenesis, immune evasion, and master regulation of gene expression (Bunnik et al., 2018). Among these are the *var* genes, a family of 60 virulence genes responsible for the antigenic variation of the parasite and evasion of the host immune system. Interestingly, only a single *var* gene is active at a given time, the other *var* genes being maintained in a perinuclear cluster of heterochromatic telomeres (Fig. 2A) (Duffy et al., 2017). This cluster is an excellent test case to validate the ability of trans-C to uncover a group of biologically important genes that co-localize in 3D from Hi-C data.

**Figure 2:**
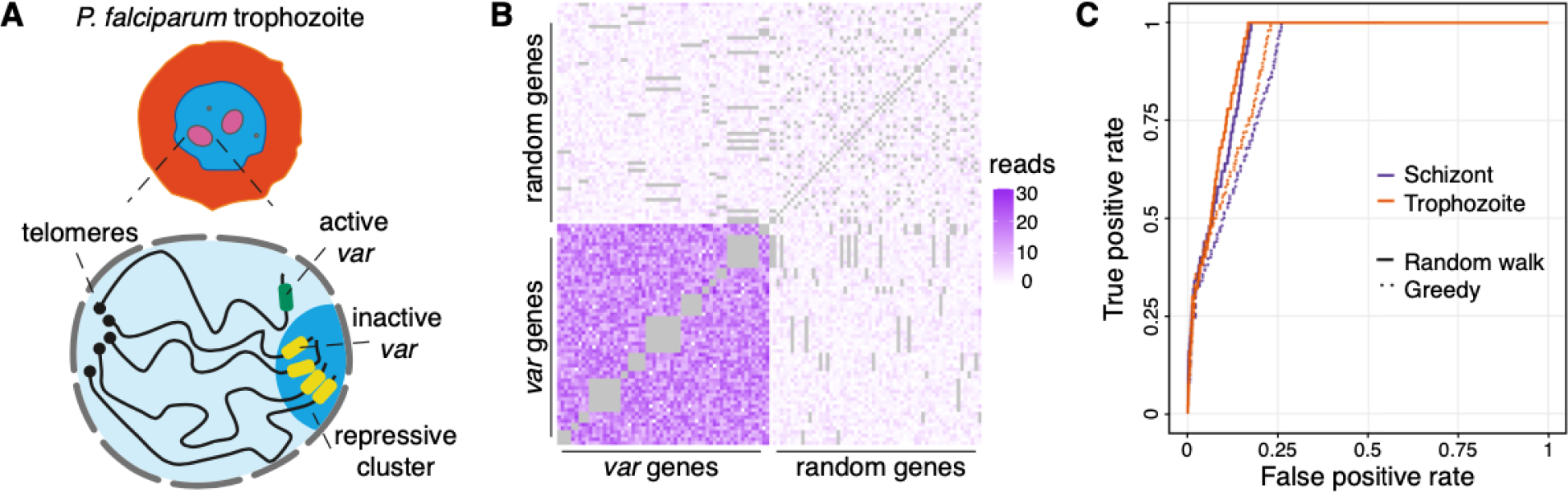
**(A)** Schematic of *Plasmodium falciparum* in the trophozoite stage of its red blood cell life cycle, with a zoomed in view of the nucleus highlighting its Rabl-like structure and the clustering of the *var* genes in a repressive heterochromatic cluster. **(B)** Contact heat map comparing *trans* interactions among all 60 *var* genes *versus* 60 randomly selected bins. *Cis* contacts are in grey. **(C)** Performance evaluation of trans-C-mediated identification of *var* gene clustering. We plot the ROC curves for two different life stages of P. falciparum. The *var* gene clustering is essential in both stages and is uncovered by the random walk algorithm of trans-C with similar high AUROC (0.94 and 0.93 for trophozoite and schizont, respectively). We also report the performance of a simpler greedy heuristic (dotted lines).

To this end, we downloaded the Hi-C data for two stages of the *P. falciparum* life cycle, trophozoite and schizont, both of which are known to be characterized by *trans* contacts between *var* genes (Ay et al., 2014). We binned the genome at 10 kb resolution and processed the data as described in detail in Methods. To visually highlight the *var* cluster, we extracted the bins containing *var* genes and also drew 60 bins at random from the full set of genomic loci. The submatrix of *trans* contacts formed by the concatenation of the two sets of bins shows a striking contrast between the *var* and non-var loci, as expected (Fig. 2B). Next, we selected three *var* genes at random to act as seed loci and examined whether trans-C (with *α* = 0.5) could automatically identify the remaining 57 *var* gene loci. For comparison we used as a baseline a method based on a greedy heuristic which iteratively selects the bins that interact most strongly with the selected loci (see “greedy heuristic” in Methods for details). For each approach, we plotted a receiver operating characteristic (ROC) curve, in which each element is a genomic bin, labeled as positive (var gene) or negative (other loci; Fig. 2C). In both *Plasmodium* life stages, trans-C quickly found the majority of the *var* genes by ranking their corresponding bins highly: of the top 50 bins, 28 contained a *var* gene, and all 60 *var* genes were recovered within the top 280 bins. trans-C outperformed the greedy heuristic baseline, with an area under the ROC curve (AUROC) of 0.94 compared to 0.88 (Fig. 2C). This demonstrates that a random walk approach is more suited to the task of identifying *trans* cliques even in the context of a remarkably clear example.

### Identification of Greek islands regulating the expression of mouse olfactory receptor genes

To further validate trans-C we turned to the mouse and its 2.6 Gb diploid nuclear genome, consisting of 40 chromosomes. Studying chromatin organization in mouse olfactory sensory neurons (mOSN), the Lomvardas lab discovered that chromatin regions associated with olfactory receptor gene clusters from 18 chromosomes make specific and robust interchromosomal contacts that increase in strength with differentiation (Lomvardas et al., 2006; Markenscoff-Papadimitriou et al., 2014; Monahan et al., 2017). These contacts are orchestrated by intergenic olfactory receptor enhancers that form a multi-chromosomal super-enhancer that drives the monoallelic and stochastic expression of a single mouse olfactory receptor gene (Fig. 3A) (Monahan et al., 2019). The mOSN-specific *trans* contacts are arguably the strongest *trans* contacts in a mammalian genome known to date; the regions involved in such interactions were dubbed “Greek islands” since they are sprinkled across the chromosomes as the tiny islands are in the Mediterranean sea. Importantly, in horizontal basal cells (HBCs), the quiescent stem cell progenitors of mOSNs, these inter-chromosomal contacts are absent.

**Figure 3:**
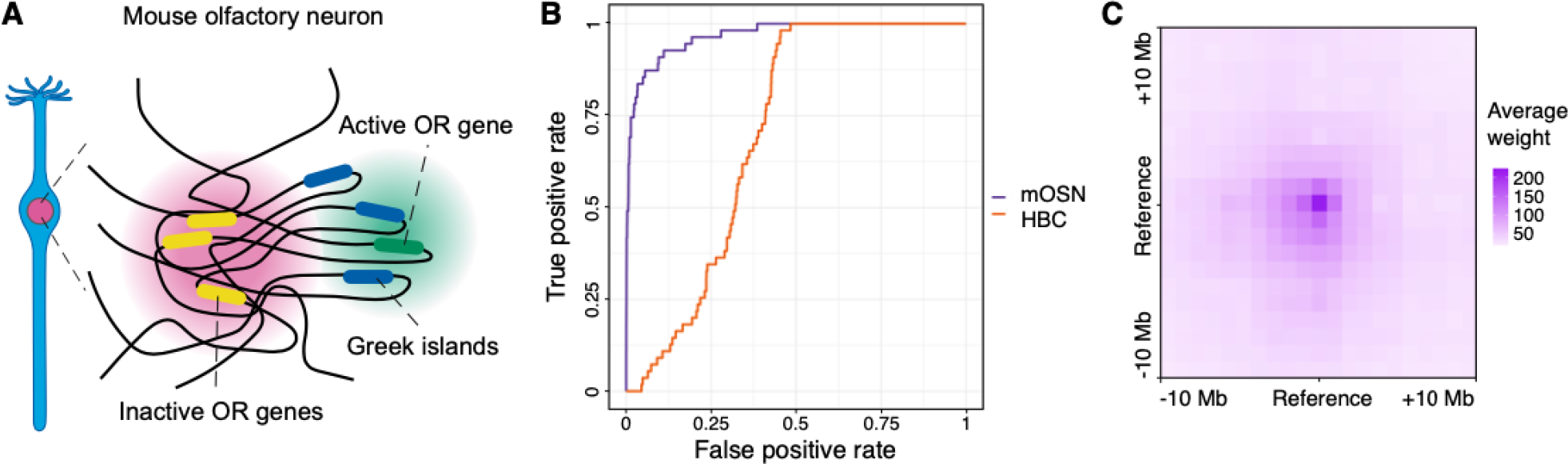
**(A)** Schematic of inter-chromosomal contacts in mouse olfactory neurons (mOSN). The Greek islands form a multi-enhancer hub that is segregated from the inactive olfactory receptor (OR) genes. **(B)** Performance evaluation of trans-C-mediated identification of Greek islands clustering. We plot the ROC curve for *α* = 0.5 in mOSNs versus their progenitors (horizontal basal cells, HBCs). Trans-C correctly identifies Greek islands clustering specifically in mOSNs. **(C)** Aggregated heatmap of *trans* contacts among the top 60 loci selected by trans-C in mOSN. Each square in the grid represents an average 250 kb bin in a Hi-C matrix of 41 *×* 41 bins centered at each interacting pair of loci (reference). The exhibited spot-like structure highlights the highly specific nature of the inter-chromosomal interactions of the Greek islands.

We applied trans-C to a published mOSN Hi-C data (Monahan et al., 2019) using the same resolution as the original analysis, 250 kb. From the list of 63 previously reported Greek islands, we randomly selected five to use as seeds, and we measured the ability of trans-C to uncover the remaining 58. As a negative control, we used the HBC Hi-C data. As expected, trans-C successfully identifies the Greek islands in mOSNs (AUROC = 0.93; Table S1), while it fails to do so effectively in HBCs (AUROC = 0.71, Fig. 3B). At a false positive rate of 10%, 95% of known Greek islands are identified, though we speculate that some of the false positives may actually represent previously unidentified Greek islands.

To visually verify whether trans-C detects specific interchromosomal contacts, we selected the top 60 predicted bins from the ranked stationary distribution (30% of which are known Greek islands). For each pair of loci from this set of 60, we extracted from the Hi-C data a 41 by 41 matrix centered at their interaction, and we averaged these matrices (Fig. 3C). The resulting contact heatmap exhibits very strong punctuated signal in the middle, suggesting that the top 60 loci ranked by trans-C form specific interactions that are not driven by larger, non-specific “neighborhood” features. Thus, trans-C efficiently pinpoints *trans* cliques even in a complex eukaryotic genome.

We also used the Greek island data set to evaluate how strongly the behavior of trans-C depends on its primary parameter, the random walk restart probability *α*. We varied *α* between 0 (no restart) to 1 (restart after every step) in small increments. We observed that the performance of trans-C is stable in the range [0.3, 0.7], while it deteriorates significantly in the two extremes when it approaches 0 or 1 (Fig. S1). This behavior is expected: when *α* is close to 0 the random walk restarts infrequently and so its stationary distribution becomes less dependent on the seeding bins and is mostly determined by the topology of the Hi-C graph. At the extreme, when *α* = 0 the walk is “memoryless” and entirely independent of the starting Greek islands (not shown). On the other hand, when *α* is close to 1 there is little or no exploration along the graph. In this setting, the Hi-C data is essentially ignored, and consequently no discoveries can be made.

### Dissecting the RBM20 splicing factory during cardiomyocyte differentiation

We next sought to explore the sensitivity of trans-C in a more challenging model in the human genome. We previously identified (Bertero et al., 2019) a network of gene loci that increase their association interchromosomally during cardiac development of human pluripotent stem cells (hPSCs) and are targets of the muscle-specific splicing factor RBM20 (Fig. 4A). Functional experiments indicated that the main RBM20 target, the large *TTN* pre-mRNA (which contains over 100 binding sites for RBM20), nucleates RBM20 foci that promote *trans* interactions with secondary RBM20 targets, which maximizes the efficiency of their alternative splicing. We therefore dubbed the network a “trans interacting chromatin domain” (TID) and the resulting structure a “splicing factory”. Of note, however, the cumulative interaction score of the TID calculated from shallow Hi-C data (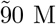 contacts) was only modestly enriched compared to a null model (p = 0.05), indicating that these interactions are less easily detected by Hi-C and are likely to be much more transient in nature compared to those involving the Greek islands.

**Figure 4:**
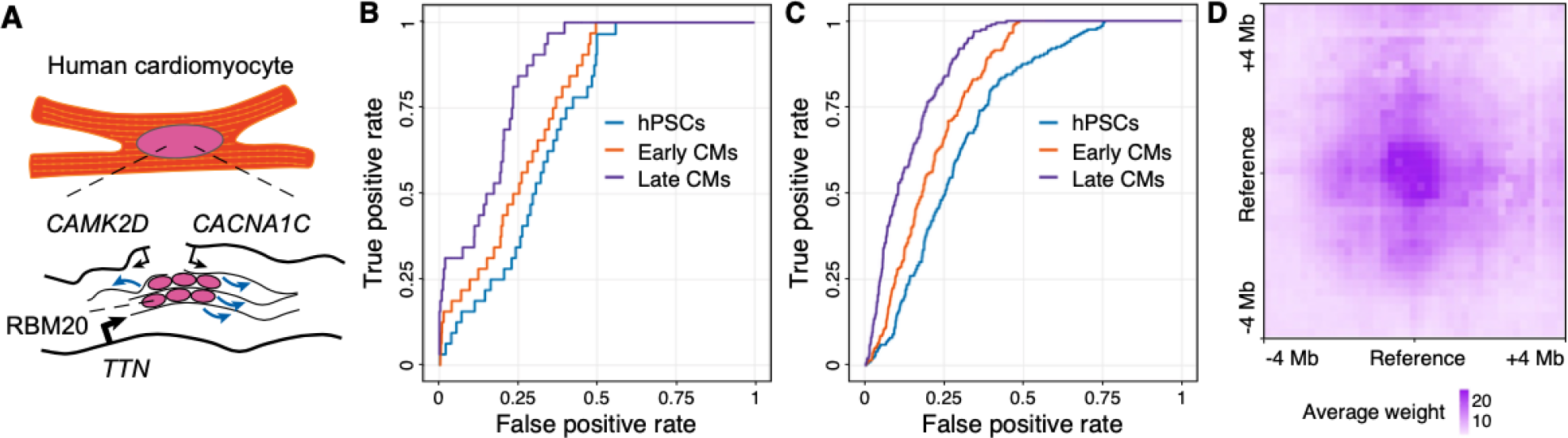
**(A)** Schematic of the RBM20 splicing factory, a muscle-specific inter-chromosomal structure organized by the *TTN* pre-mRNA, which binds to *>* 100 copies of RBM20, thus nucleating foci that engage with other RBM20 targets to promote their alternative splicing (blue arrows). **(B)** Performance evaluation of trans-C in uncovering the RBM20 splicing factory in early (day 15) *versus* late (day 80) cardiomyocytes (CMs) differentiated from pluripotent stem cells (hPSC; also analyzed as “day 0” baseline control). ROC curves are calculated using a list of *TTN* -interacting from our previous Hi-C study (Bertero et al., 2019). **(C)** Similar to B, but the positive list consists of loci directly bound by RBM20. **(D)** Aggregated heatmap of *trans* contacts between the top 60 loci selected by trans-C in late CMs. Each square in the grid represents an average 100 kb bin in a Hi-C matrix of 41 *×* 41 bins centered at each interacting pair of loci extracted from the Hi-C data (reference). The dense region in the middle reveals the specific nature of the *trans* interactions at the RBM20 splicing factory.

We set out to test whether trans-C would re-identify the RBM20 TID in an independent, more deeply sequenced Hi-C dataset of hPSC differentiation into the cardiac lineage (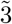 billion read pairs per sample (Zhang et al., 2019)). Besides various progenitors and early hPSC-derived cardiomyocytes (hPSC-CMs), this dataset also contains older hPSC-CMs obtained after 80 days of in vitro differentiation and FACS-sorted using an expression reporter for the mature marker ventricular myosin light chain 2 (MLC-2v; *MYL2* gene). We first attempted to recover the *trans* network of 16 RBM20 target genes from our original report, using five of them (*TTN, CACNA1C, CAMK2D, KCNIP2, CAMK2G*) as seeds for trans-C. Figure 4B shows the ROC curve for day 0 (hPSCs), 15 (early CMs) and 80 (late CMs). The best performance is achieved using Hi-C data from day 80 (AUROC = 0.84; Table S2), when cardiac development is most advanced, in line with the important role of RBM20 in cardiac maturation (Guo et al., 2012). Second is day 15 (AUROC = 0.78), and last is day 0 (AUROC = 0.75). The improvement in ROC area as differentiation advances is in line with biological expectation (i.e., RBM20 is not expressed at day 0, moderately expressed at day 15, and maximally expressed at day 80). We note, however, that the performance at day 0 is better than random, suggesting that some structure that brings the loci close together is present even in undifferentiated cells.

Encouraged by these results, we decided to use trans-C to expand our knowledge of the RBM20 TID. Our original list of 16 genes was not the result of an unbiased search but rather reflected our prior knowledge of RBM20 biology: these 16 genes were the known splicing targets of RBM20 in both human and rat hearts that were also upregulated in hPSC-CM. As an alternative strategy to identify genes involved in the RBM20 TID in an unbiased fashion, we hypothesized that such genes would encode for transcripts most strongly bound by RBM20 and thus enriched within the splicing factory. To test this hypothesis we leveraged our recent dataset that measured RBM20 binding to RNAs using enhanced UV crosslinking and immunoprecipitation (eCLIP) (Van Nostrand et al., 2016).

We downloaded RBM20 eCLIP data from hPSC-CMs (Fenix et al., 2021), binned the genome at the same resolution as the Hi-C matrix, 100 kb, and counted the number of peaks that fall in each bin. We selected the five bins with the most eCLIP peaks, which contained the genes *TTN, SLC8A1, OBSCN, NEAT1*, and *LBD3*. Using these as seed loci, we ran trans-C with *α* = 0.5 on Hi-C matrices from differentiating hPSC-CMs (Zhang et al., 2019). Our goal was to test whether trans-C would uncover the remaining 202 bins with eCLIP peaks. We note that this experimental setup is very different from the previous ones. Here, we are using Hi-C data to find binding sites in an orthogonal eCLIP dataset. The resulting ROC curves (Fig. 4C) show the same trend discussed for Figure 4B: best performance is at day 80 (Table S3), second at day 15, and last at day 0, consistent with biological expectation.

Next, we performed a second analysis in which we restricted the list of RBM20 targets to those whose RNA is bound by RBM20 on at least three sites and is differentially spliced in hPSC-CMs with RBM20 knocked out (Fenix et al., 2021). The resulting list of 45 high confidence RBM20 targets was efficiently recovered by trans-C in day 80 hPSC-CMs, with AUROC = 0.84 (Fig. S2), compared to AUROC of 0.82 for the full list of RBM20 bound loci.

Lastly, similarly to our observation for *trans* contacts between the Greek islands (Fig. 3C), the aggregated contract frequency heatmap for loci involved in the RBM20 splicing factory showed a clear punctuated pattern, supporting the spatial specificity of these interactions (Fig. 4D). In all, we conclude that trans-C captures even weak and/or unstable yet biologically meaningful *trans* subnetworks associated with RNA biogenesis and, further, that our method can be used in conjunction with independent datasets to interrogate different but spatially proximal processes.

### TF binding loci exhibit strong interactions in trans

The identification of dense subnetworks of *trans* Hi-C contacts that are enriched for RBM20 targets supports our hypothesis that RNA biogenesis influences 3D chromatin organization by bringing into proximity co-regulated nucleic acids, so as to maximize the efficacy and specificity of their processing (Fig. 5A (Bertero, 2021)). We specifically propose that, as in the case of RBM20, RNA factories arise from the clustered binding of *trans*-acting factors to one or more core co-regulated genes and/or their encoded transcripts, which in turn recruit accessory targets of the same factors. This hypothesis predicts the existence of both transcription factories specialized for certain TFs and other RNA factories specialized for various RNA binding and regulatory proteins. We set out to test this hypothesis systematically using trans-C, as an example of its potential applicability to address biological questions. First, we focused on TFs, hypothesizing that the genes most strongly bound by a given TF would be associated with strong TIDs. To test this notion, we used the most deeply sequenced Hi-C dataset to date: an ultra-deep Hi-C map of human GM12878 lymphoblastoid cells (Gu et al., 2021), which contains in total a staggering 33 billion mapped reads, 3.7 billion of which correspond to *trans* contacts. We performed our analysis at 100 kb resolution, which results in non-zero counts for 85% of all pairwise *trans* contacts. The ENCODE project (ENCODE Project Consortium, 2012) produced chromatin immunoprecipitation sequencing (ChIP-seq) data for 110 TFs in this cell line, providing an ample resource to test our hypothesis in a systematic manner. For each ChIP-seq dataset we subdivided the human genome at 100 kb resolution and counted the number of peaks in each bin.

**Figure 5:**
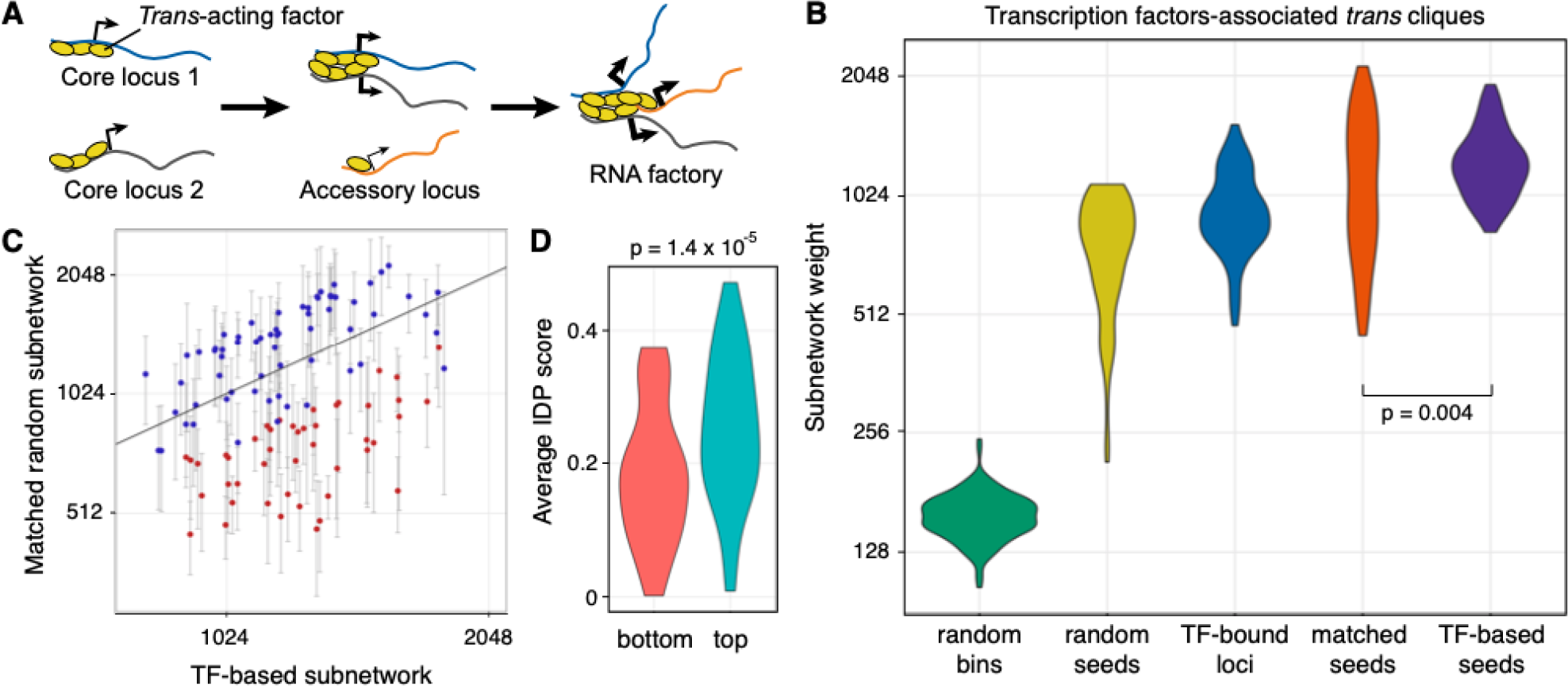
**(A)** Schematic of the mechanistic hypothesis for the formation of specialized RNA factories involving *trans* interacting chromatin domains. Multiple copies of *trans*-acting regulatory factors (i.e., transcription or splicing factors) bind to core nucleic acids, aggregate to form new clusters and/or enrich pre-existing ones, and recruit accessory co-regulated nucleic acids. RNA factories promote the efficacy and accuracy of RNA biogenesis processes (thicker black arrows). **(B)** Trans-C-identified subnetworks in lymphoblastoid cells built from loci characterized by strong binding of transcription factor (TF) have dense contacts. We plot the distribution of subnetwork weights for five types of equally sized sets of loci: (1 - green) sets of randomly drawn loci; (2 - yellow) sets built by trans-C from a seed of five randomly drawn loci; (3 - blue) sets of loci with the highest number of ChiP-seq peaks for a given TF; (4 - orange) sets built by trans-C from a random seed of five loci whose starting subnetwork weight was matched to the seed of group 5; and (5 - purple) sets built by trans-C from a seed of five loci with highest number of ChiP-seq peaks for a given TF. On average, sets seeded from loci most strongly bound by TFs interact more strongly in *trans* than any of the other four types of sets of loci, including the stringent “matched seeds” controls (p-value by Mann-Whitney test). **(C)** For each TF analyzed in B, we compare the weights of subnetworks obtained with trans-C from “TF-based seeds” (single data point) and “matched seeds” (average of 20 subnetworks ± standard deviation). In red are comparisons with significantly different weights (p-value *<* 0.05 after FDR correction). **(D)** The strongest TF-based subnetworks correspond to TFs with a higher intrinsically disordered protein (IDP) score. We plot the IDP scores for the bottom and top quartile of TFs ranked by the weight of the resulting trans-C “TF-based seeds” subnetwork (purple violin plot in B). The difference is statistically significant by Mann-Whitney test.

First, we took the 40 bins with the most peaks for each TF and calculated the weight of the subnetworks formed by these bins. The distribution of this subnetwork weight across all 110 TFs is shown in blue in Figure 5B. For comparison, we randomly drew 1000 sets of 40 bins and plotted the distribution of their weight in green. Clearly, the subnetworks of loci selected based on ChIP-seq peak density form stronger interactions in *trans* than random sets of loci. This is a first important hint that many TFs may be indeed involved in specific *trans* contact networks.

Second, for each TF individually, we formed a seed by selecting the five bins with the most peaks from its corresponding ChIP-seq track, and we ran trans-C to identify a set of potential interactors in *trans*. We took the top 40 predicted bins for a given TF and observed that these bins are enriched with ChIP peaks not only for the TF that spawned the seed, but also for ChIP peaks of other TFs (Fig. S3). This is not surprising because many TFs act in concert, and many loci contain proximal binding sites of several TFs (Ibarra et al., 2020). When examining the weights of subnetworks formed by trans-C (TF-based seed, purple in Fig. 5B) we noted that they were heavier on average than the subnetworks based on the ChIP-seq signal only (TF loci, blue). This observation validates that trans-C finds loci that interact even more strongly in *trans* with the seed bins than the just the bins most bound by the respective TF.

We also assessed how well trans-C can build dense subnetworks when it is seeded from biologically unrelated loci. To that end, we first drew 20 times five random loci to use as seeds and ran trans-C. The subnetworks it built were significantly weaker than the TF-based ones. This is likely due to the fact that a randomly drawn seed set likely includes loci that are not interacting with one another, while the loci in the TF-based seed tend to have strong interactions in *trans*. Thus, in order to establish a more stringent baseline, for each TF we repeatedly drew five random bins until we found a set that has the same total weight as the TF-based seed. We called this random seed “matched random” and generated 20 such sets for each TF. Interestingly, when using this matched random seed the subnetworks that trans-C found were once again weaker on average than the ones it found using TF-based seed (Mann-Whitney test *p* = 0.004; orange in Fig. 5B). At an individual level, the weight of subnetworks for 40% (44 out of 110) TFs identified from the TF-based seeds were significantly stronger than those from matched random seeds (Fig. 5C) These observations confirm the common sense conception that the interchromosomal interactions of biologically unrelated loci are mostly noise, while providing more rigorous support to the hypothesis that co-regulated loci are often enriched for *trans* contacts.

Examining the distribution of the trans-C subnetwork weights identified for different TFs (Fig. 5B, purple) we noticed a bimodal distribution, indicating that some TFs are associated with stronger TIDs. This bimodality did not correlate with differences in the expression level of the two groups of TFs (data not shown), nor in their preference to bind to loci in the A or B compartments (Fig. S4). Intrinsically disordered regions (IDRs) within proteins, which lack a defined tertiary structure and are thus prone to self-aggregation, are emerging as an important mediator of subcellular condensates involved in multiple aspects of cell function, including nuclear regulations (Wright and Dyson, 2015). We thus investigated the correlation between the intrinsic disorder in TF structure and the strength of the trans-C subnetworks they are associated with. For this analysis, we took the TFs whose seeds gave rise to the strongest and weakest subnetworks (top and bottom 25% of the purple distribution in Fig. 5B, Table S4). We used the tool developed by the Dosztányi lab (Mészáros et al., 2018) to calculate an average intrinsically disordered protein (IDP) score for each TF in the two groups, and plotted them in Figure 5D. Interestingly, the difference between the two groups is statistically significant (Mann-Whitney test *p* = 1.4 × 10^*−*^5), suggesting that the TFs with more intrinsically disordered regions form stronger interactions in *trans*. In all, trans-C allowed us to identify a large set of TF enriched for IDR regions that are involved in strong TIDs and that may thus be important in efficient transcriptional regulation of their target genes.

### RNA binding proteins create strong *trans* interacting sets

Encouraged by these results on TF subnetworks, we also examined whether RNA binding proteins (RBPs) are generally associated with TIDs. Because only a few RBP binding profiles are available for GM12878, for this analysis we turned to human K562 cells, another lymphoblastic line, for which ENCODE reports 139 eCLIP datasets. Accordingly, we used a Hi-C matrix for that cell line from the ENCODE portal, although it had far fewer *trans* reads (360 million) than the deeply sequenced GM12878. Similar to our TF analysis, we first divided the genome into 100 kb bins and counted the number of peaks in each bin for each RBP. Then, for each RBP individually, we formed a seed by selecting the five bins with the most peaks and ran trans-C to identify a set of potential interactors in *trans*. To form a null model per RBP, we repeatedly drew 20 contact frequency-matched random seeds. We report the average total weight of the matched random seeds compared to the RBP seed in Figure 6.

**Figure 6:**
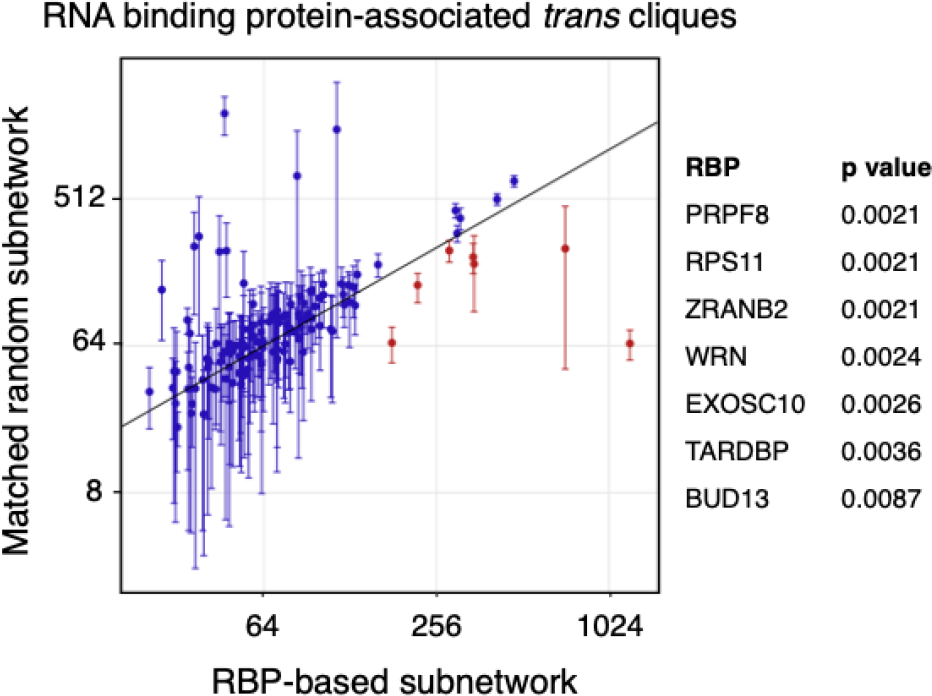
A subset of trans-C-identified subnetworks in lymphoblastoid cells built from loci characterized by strong binding of RNA binding protein (RBP) have denser contacts than the corresponding random null model. We plot the weight of a RBP-based subnetwork and the average weight of 20 matched random seed subnetworks (error bars correspond to the standard deviation) for 139 RBPs. In red and reported on the right are those with significant p-values after FDR correction.

Most RBP subnetworks built by trans-C were comparatively as dense as those from the corresponding matched random seed, lying broadly along the *y* = *x* line. Nevertheless, several outliers are notably denser. To assess this observation quantitatively we performed a signed ranked test per RBP and FDR controlled the corresponding p-values. Seven proteins had corrected p-values lower than 0.05, which we considered as a significance threshold (indicated in red in Fig. 6 and listed in Table S5). Distinctly from TFs, RBPs associated with significantly stronger trans-C subnetworks were not characterized by higher IDP scores (data not shown), indicating that other characteristics may explain their specific behavior in *trans* genome organization. In all these analyses indicated that, unlike TFs, RBPs are not generally involved in stronger than average TIDs, but that a small subset of RBPs is associated with very strong ones (enriched up to *∼* 16 fold).

## Discussion

Potential inter-chromosomal interactions occupy 90% of the pairwise 3D DNA contact space, and a sizable fraction of measured interactions. While in certain species whose nucleus is characterized by chromosome territories — such as humans and other mammals — a large fraction of *trans* contacts are likely nonspecific, illuminating an even small fraction of specific and functional inter-chromosomal interactions may provide important advances in our understanding of nuclear mechanisms such as transcription and splicing. In this context, trans-C is an important step towards refined analytical methods to probe the *trans* contact space for functional gene networks.

The study of *trans* contacts requires statistical methods designed for the specific task at hand. Approaches devised for the analysis of *cis* interactions control for some biases that are not applicable to *trans* ones, such as correction for the linear genomic distance between the interacting loci. To date, most analytical tools for Hi-C data have been limited to *cis* interactions (Lin et al., 2019). Recent network-based strategies to study inter-chromosomal interactions from bulk and single cell Hi-C (Kaufmann et al., 2015; Bulathsinghalage and Liu, 2020; Joo et al., 2023) proposed probabilistic models that focus on identifying large patterns of *trans* contacts (i.e., those involved in nuclear speckles and nucleoli) rather than small sets of interactions linked to a specific process. Trans-C is, to the best of our knowledge, the first method that controls for chromosome territory biases to identify gene networks that “stand out” from other *trans* interactions resulting from the random intermixing of neighboring chromatin domains.

We first validate the ability of trans-C to detect known examples of functional *trans* contacts. We find that the approach outperforms a simpler greedy heuristic even in the case of the simple genome of P. falciparum, which is characterized by remarkable *trans* contacts among *var* genes. In more complex and larger mammalian genomes, trans-C not only identifies with high precision the mOSN Greek islands, but also the less striking example of *trans* contacts represented by the RBM20 splicing factory. Thus, trans-C may find applicability across nuclear genomes with different sizes and types of organization, and *trans* contacts of varying strength. We demonstrate the utility of trans-C by using it to systematically seek for *trans* cliques around loci most strongly bound by one of many TFs or RBPs. These analyses support the existence of a large number of statistically significant TIDs readily measurable from Hi-C data, particularly in the case of intrinsically disordered TFs. The concept of “bookmarked” transcription factories, Pol II clusters that are specifically enriched for a set of TFs and their target loci, has been proposed over a decade ago (Cook, 2010), but examples of this concept have been sparse (reviewed in (Bertero, 2021)). Our analysis of 110 TFs provides an important piece of evidence to support this model for a sizable fraction of TFs, though a mechanistic dissection of these leads will be required to firm up the conclusions.

Intriguingly, the few RBPs associated with significant TIDs are involved in a wide variety of functions. Not only do we identify several factors involved in major and minor spliceosomes (PRPF8 and BUD13), but we also identify alternative splicing regulators (ZRANB2), a multifunctional RNA processing factor (TARDBP), a component of the RNA exosome complex (EXOSC10), a ribosomal protein (RPS11), and even a DNA helicase involved in homologous recombination (WRN). We speculate that these factors exemplify a wide range of chromatin structures involving both *cis* and *trans* interactions that regulate not only transcription but also other aspects of nucleic acid biology such as DNA replication and repair, or various aspects of RNA biogenesis. Notably, several of the RBPs highlighted by our trans-C analysis are known to be mutated in severe human monogenic diseases: PRPF8 in retinitis pigmentosa (McKie et al., 2001), WRN in Werner syndrome (Yu et al., 1996), and TARDBP in amyotrophic lateral sclerosis (Sreedharan et al., 2008). Moreover, mutations in ZRANB2 have been linked to unfavorable prognosis in breast and liver cancer (Tanaka et al., 2020), while RPS11 has been shown to be a key player in poor outcomes of glioblastoma patients (Dolezal et al., 2018). Whether misorganization of *trans* genome architecture is implicated in the pathogenesis of these diseases is an interesting topic for future investigations.

Overall, our work focuses on poorly studied between-chromosome contacts and provides an efficient computational framework for identifying potentially biologically important loci that interact in *trans*. We demonstrate the flexibility and sensitivity of trans-C and provide examples of how our approach can be used to identify candidate gene networks for subsequent hypothesis-driven studies. Application of trans-C to the growing number of Hi-C datasets from the ENCODE (ENCODE Project Consortium, 2012) and 4D Nucleome consortia (Reiff et al., 2022) will reveal novel cell- or disease state-specific *trans* networks. Minor adaptations of trans-C will also allow exploration of other proximity-ligation independent assays, such as SPRITE, GAM, and TSA-seq (Beagrie et al., 2017; Chen et al., 2018; Quinodoz et al., 2018), which will collectively offer the potential to accurately characterize inter-chromosomal architecture at varying spatial resolutions.

## Methods

### Overview

Trans-C takes as input a Hi-C contact matrix *H* of interaction counts and an initial set *S* of loci of interest (“seed loci”) and outputs a set of loci *U* (containing *S*) that interact strongly together in *trans*. We refer to *U* and its associated edges as a “dense subgraph.” We model the Hi-C interaction matrix *H* as a weighted graph *G* = (*V, E, W*) with nodes *V* corresponding to the genomic loci (bins), edges *E* between pairs of loci, and weights *W* on the edges reflecting the strength of the interactions represented by the edges. Thus, the weight *w*_*ij*_ on edge *e*_*ij*_ between loci *i* and *j* corresponds to the contact matrix entry *h*_*ij*_. Our goal is to find a subset of loci that exhibit strong inter-chromosomal contacts. We propose two methods to solve this problem, one that uses a random walk with restarts and a second that formulates the problem as a dense subgraph optimization and solves it using a fast greedy heuristic.

**Table 1:**
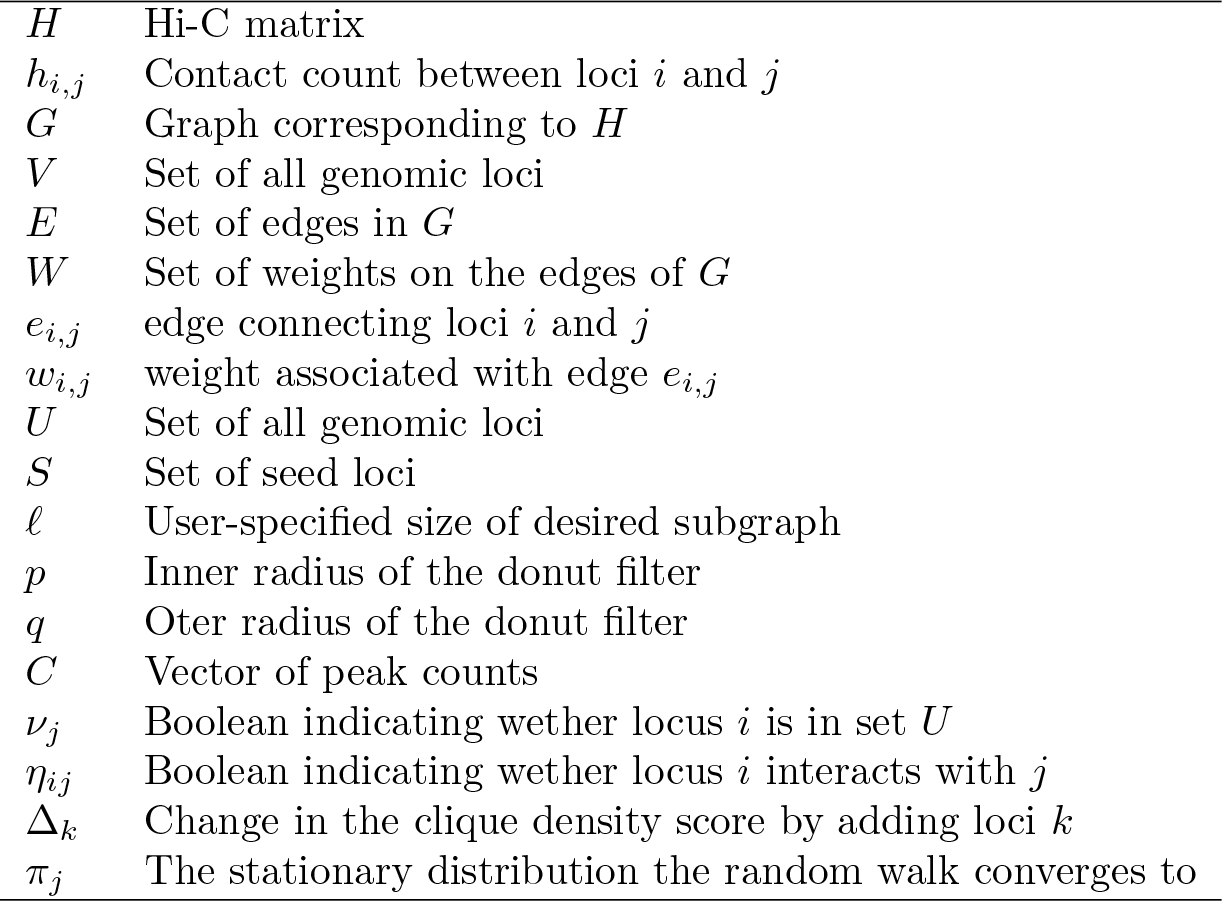
Notation.

|  |  |
| --- | --- |
| $H$ | Hi-C matrix |
| $h_{i,j}$ | Contact count between loci $i$ and $j$ |
| $G$ | Graph corresponding to $H$ |
| $V$ | Set of all genomic loci |
| $E$ | Set of edges in $G$ |
| $W$ | Set of weights on the edges of $G$ |
| $e_{i,j}$ | edge connecting loci $i$ and $j$ |
| $w_{i,j}$ | weight associated with edge $e_{i,j}$ |
| $U$ | Set of all genomic loci |
| $S$ | Set of seed loci |
| $\ell$ | User-specified size of desired subgraph |
| $p$ | Inner radius of the donut filter |
| $q$ | Oter radius of the donut filter |
| $C$ | Vector of peak counts |
| $\nu_j$ | Boolean indicating wether locus $i$ is in set $U$ |
| $\eta_{ij}$ | Boolean indicating wether locus $i$ interacts with $j$ |
| $\Delta_k$ | Change in the clique density score by adding loci $k$ |
| $\pi_j$ | The stationary distribution the random walk converges to |

### Random walk with restarts

Our first solution carries out random walks with restarts over the graph *G* and then uses the results of the random walk to select the nodes in *U* . The walk is initiated from the set of seed loci *S*. At each step, with probability *α* the walk restarts from a randomly selected seed locus, and with probability 1 *− α* the walk moves to a neighboring locus picked probabilistically based upon *W* . Specifically, if 𝒩 (*i*) are the loci *i* interacts with, then the walk goes from locus *i* to locus *j ∈* 𝒩 (*i*) with probability proportional to *w*_*i,j*_*/*Σ_*k∈𝒩* (*i*)_ *w*_*i,k*_. That is, if at time *t* the walk is at locus *i*, then the probability that it transitions to locus *j* at time *t* + 1 is

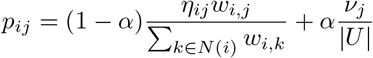

where *η*_*ij*_ = 1 if *j ∈* 𝒩 (*i*) and 0 otherwise, and *ν*_*j*_ = 1 if *j ∈ U* and 0 otherwise. Hence, the guided random walk is fully described by a stochastic transition matrix *P* with entries *p*_*ij*_. This stochastic matrix is non-negative and by the Perron-Frobenius theorem it has a right eigenvector *π* corresponding to eigenvalue 1. Therefore, *πP*^*t*^ = *π*, and we can efficiently compute the stationary distribution *π* that the guided random walk converges to. The score of each locus *j* is given by the *j*th element of *π*. The loci that have high scores are most frequently visited and, therefore, are more likely relevant as they are strongly connected to the seed loci. We use these scores to rank all loci and include the top 𝓁 loci in *U*, where 𝓁 is a user-specified parameter. In this work, we use 𝓁 = 40 unless otherwise stated.

### Greedy heuristic

Our second solution builds a ranked list of loci in *U* in a greedy fashion. In this approach, we formalize our goal as finding *U ∈ V* such that the subgraph induced by |*U* | is dense; i.e., the following quantity is large:

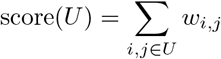

We note that when we constrain the size of *U*, the problem is computationally hard as it can be reduced to the maximum clique problem, which is NP-complete. As in the previous approach, we assume that the user specifies an initial set *S* of seed loci, as well as the desired subgraph size 𝓁. Thus, formally, the optimization problem we aim to solve is

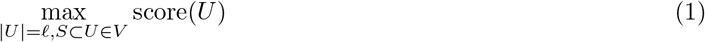

We propose to maximize Equation 1 using a greedy algorithm. The procedure begins by adding the seed loci *S* to the initially empty set *U* . Then, in each step the heuristic expands *U* by examining all loci not currently in *U* and selecting to add to *U* the one that yields the largest increase in Equation 1. Mathematically, the greedy step finds

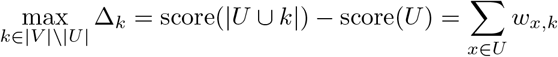

Ties are broken randomly. The greedy selection proceeds as long as Δ_*k*_ *>* 0 and |*U* | *<* 40. In practice, in the calculation of Δ_*k*_ we exclude the single strongest interaction between *k* and *U* . We do so because we don’t want a single very large *w*_*k,x*_ to dominate Δ_*k*_; instead, our aim is that all loci in *U* interact strongly.

### Data pre-processing

Prior to searching a given Hi-C matrix for dense subgraphs, we perform three pre-processing steps. First, we normalize the Hi-C matrix using the iteractive correction and eigenvalue decomposition (ICE) procedure (Imakaev et al., 2012). This procedure iteratively normalizes rows and columns of the matrix, producing as output a matrix in which the marginal values are all equal to a specified constant. We carry out the procedure on the entire Hi-C matrix, including *cis* and *trans* contacts, using the python package “iced” (Servant et al., 2015).

Second, we adjust the matrix entries to account for the fact that chromosomes tend to occupy specific regions of the nucleus, called *chromosomal territories*, and as a result some pair of chromosomes interact more frequently. For each pair of chromosomes *L* and *M* (*L* ≠ *M*) we find the total number of interactions between any locus *i* in *L* and *j* in *M* : *T*_*L,M*_ = Σ_*∀i∈L*;*∀j∈M*_ *h*_*ij*_. If *T* is the total number of trans-interactions in *H*, then we rescale contact count *h*_*ij*_ as 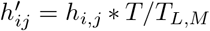. During this step, we set all *cis* contacts (*i* and *j* are on the same chromosome) to zero.

Third, we process *H* using a “donut filter” to emphasize points that are local minima in the 2D contact map (Rao et al., 2014). Given a *trans* contact (*i, j*), we define its donut background as the set of all loci that are at least *p* loci away from (*i, j*) but no further than *q* loci away and which do not lie along the *i* or *j* axes. Intuitively, *p* is the radius of the hole of the donut centered at (*i, j*), *q* is the outer radius of the donut, and the donut has been sliced in four pieces along the *i* and *j* axes. Mathematically,

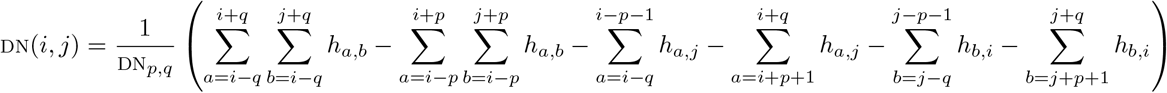

where we divide the sum by the total number of loci DN_*p,q*_ in the donut to obtain the average strength of interactions in the donut. The enrichment for the contact (*i, j*) with respect to its local background can then be calculated as *h*_*i,j*_*/*DN(*i, j*). In practice, we select *p* = 2 and *q* = 20, and we set 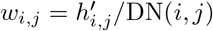.

### Datasets

We apply our method to Hi-C data from three organisms. First, for *Plasmodium falciparum*, we obtained Hi-C matrices from the trophozoite and schizont stages of the development of the parasite (accession GSE126074) (Bunnik et al., 2018). As seed loci, we selected five genes at random from a curated list of virulence genes, which are essential for the evasion of the host immune system and are known to come in close contact in the nucleus (Gardner et al., 2002).

Second, for we downloaded mouse Hi-C data from the mature olfactory sensory neuron (4DN Portal id: 4DNFI3M6726I) (Monahan et al., 2019). As seed loci, we used five randomly selected loci from the list of 63 intergenic regions labeled as Greek islands by the authors of the study.

Third, we used human Hi-C matrices from three timepoints during cardiomyocyte differentiation day 0 (4DNFIT5YVTLO), day 15 (4DNFIIOUG5RF) and day 80 (4DNFI8RH55DO) (Zhang et al., 2019). The raw data was provided at 10 kb resolution, which we converted to 100 kb resolution. From the published list of 16 RBM20 targets (Bertero et al., 2019), we selected *TTN, CACNA1C, CAMK2D, KCNIP2*, and *CAMK2G* as seed loci based on prior validation of the *trans* contacts between these loci.

In addition, we downloaded ChIP-seq data for 110 human transcription factors (TF) in the GM12878 cell line and 139 eCLIP RNA binding proteins in the K562 cell line from the ENCODE portal (ENCODE Project Consortium, 2012). We split the human linear genome in 100 kb bins, and for a given protein *t* counted the number of peaks in each bin, producing a count vector *C*_*t*_. When more than one dataset is present for a given protein, we aggregated the peaks from the datasets for that protein. For the TF analysis in the GM12878 cell line we used the ultra-deep sequenced Hi-C matrix from (Gu et al., 2021) and for the RBP analysis in the K562 cell line– the intact Hi-C matrix (ENCFF621AIY) from ENCODE.

## Declarations

### Availability of data and materials

The trans-C scripts are available in the https://github.com/Noble-Lab/trans-C repository. All of the datasets used for analyses were previously published, and their accession numbers are detailed in the relevant method sections.

## Acknowledgements

The study was funded by the National Institutes of Health (award UM1HG011531; WSN), the Giovanni Armenise-Harvard Foundation (Career Development Award 2021; AB), and the European Union (ERC, TRANS-3, 101076026; AB). Views and opinions expressed are however those of the author(s) only and do not necessarily reflect those of the European Union or the European Research Council Executive Agency. Neither the European Union nor the granting authority can be held responsible for them.

## Authors’ contributions

B.H.H. performed all of the analyses and wrote the first draft of the manuscript. W.S.N. supervised the analyses, edited the manuscript, and obtained funding. A.B. conceptualized the study, co-supervised the analyses, edited the manuscript, and obtained funding.

## Supplementary Figures

**Supplementary Figure S1:**
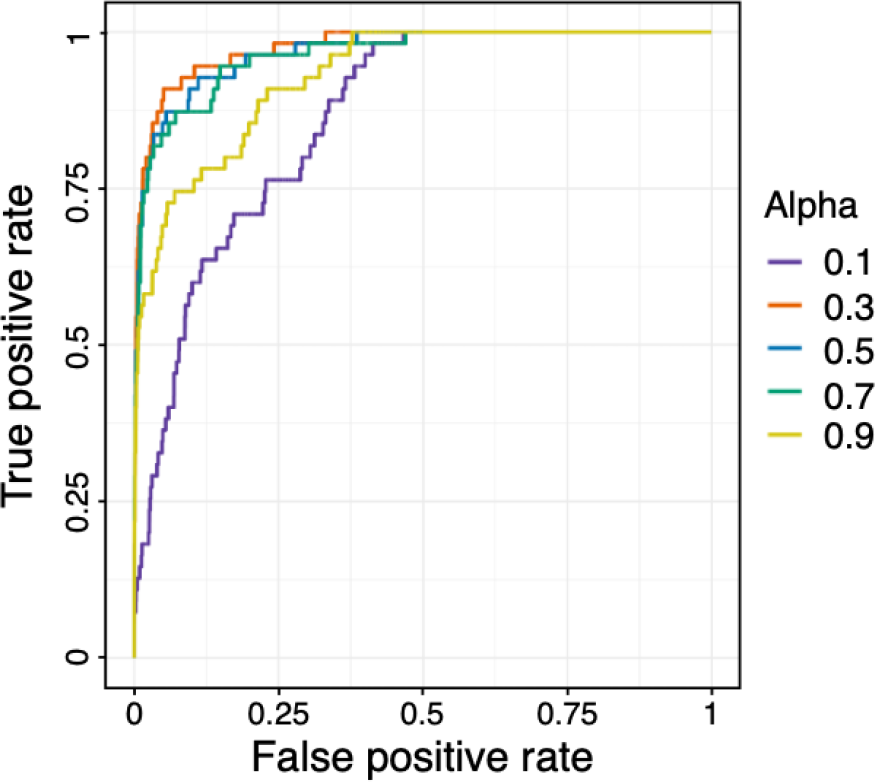
Performance of trans-C in recovering the Greek islands in mouse olfactory neurons with different values of the parameter alpha (refer to Fig. 3B).

**Supplementary Figure S2:**
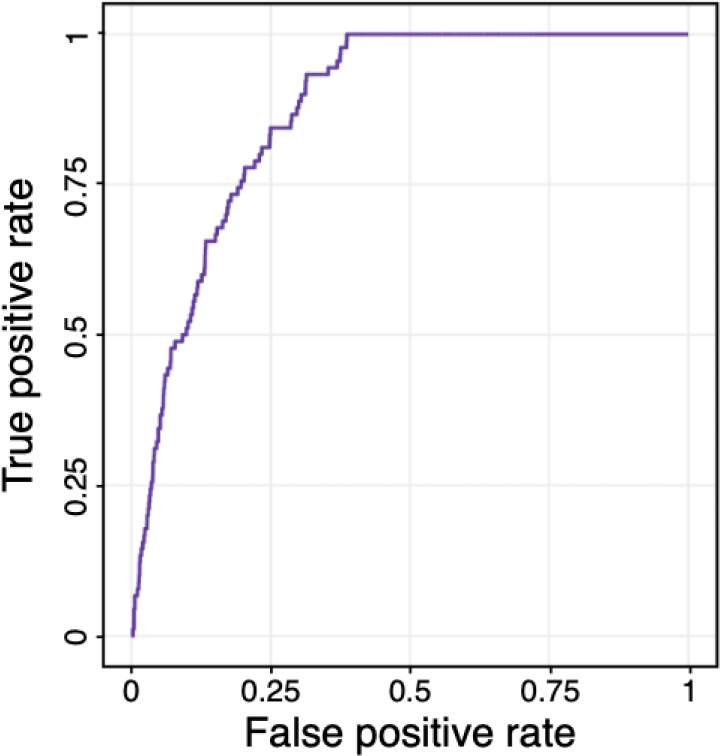
Performance of trans-C in recovering a recall list of 45 high confidence RBM20 targets in late CMs (refer to Fig. 4C).

**Supplementary Figure S3:**
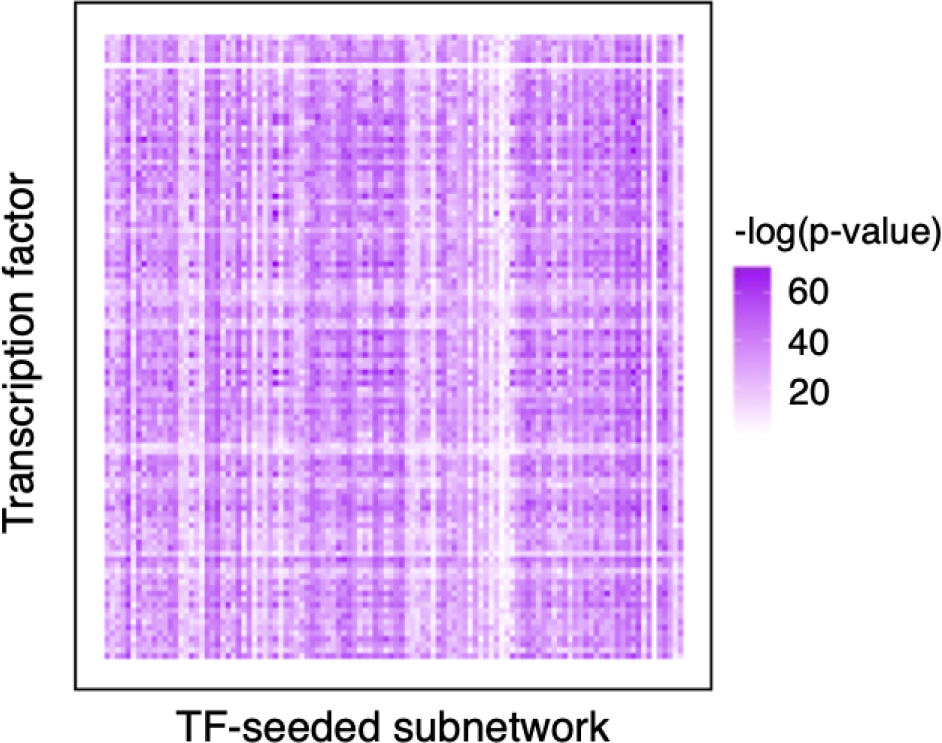
Enrichment for TF peaks in subnetworks built by trans-C from TF-based seeds (refer to Fig. 5B, violet plot). Each row corresponds to a TF and each column to a TF-based subnetwork built by trans-C. For each TF-based subnetwork we report enrichment for peaks of other TFs using a hypergeometric test, and report the negative logarithmic p-value in the corresponding cell.

**Supplementary Figure S4:**
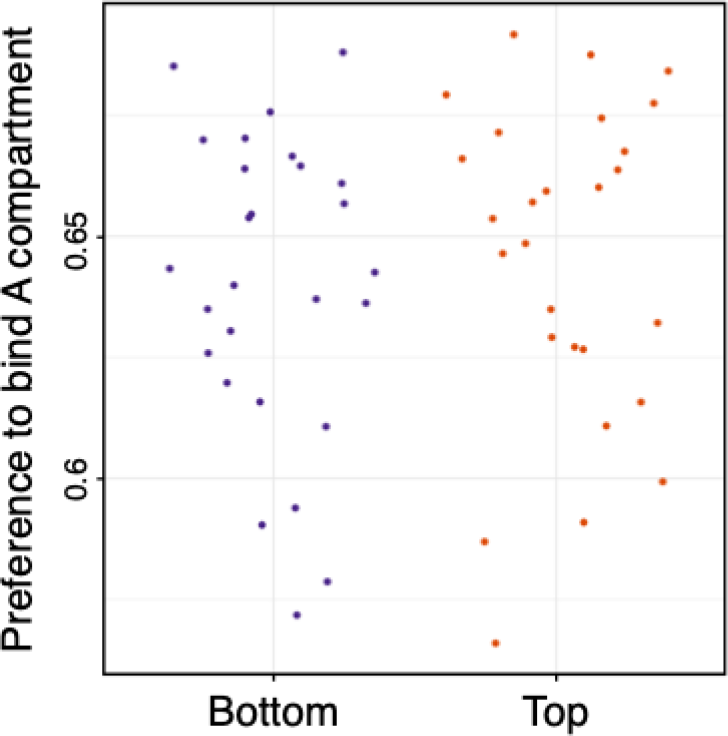
Refer to Fig. 5B, violet plot. TFs that yielded the top and bottom quantiles of subnetwork strength show no difference in their preference to bind loci in the A or B compartments. The y-axis measures the fraction of bins bound by a TF that are in A compartment.

## Supplementary Tables (online)

Supplementary Table 1: Predicted Greek islands interactors

Supplementary Table 2: RBM20 predicted interactors

Supplementary Table 3: RBM20 eCLIP predicted interactors

Supplementary Table 4: TF-based cliques p-values

Supplementary Table 5: RBP-based cliques p-values

## References

F. Ay, E. M. Bunnik, N. Varoquaux, S. M. Bol, J. Prudhomme, J.-P. Vert, W. S. Noble, and K. G. Le Roch. Three-dimensional modeling of the P. falciparum genome during the erythrocytic cycle reveals a strong connection between genome architecture and gene expression. Genome Research, 24:974–988, 2014.

R. A. Beagrie, A. Scialdone, M. Schueler, D. C. Kraemer, M. Chotalia, S. Q. Xie, M. Barbieri, I. de Santiago, L. M. Lavitas, M. R. Branco, J. Fraser, J. Dostie, L. Game, N. Dillon, P. A. Edwards, M. Nicodemi, and A. Pombo. Complex multi-enhancer contacts captured by genome architecture mapping. Nature, 543(7646):519–524, 2017.

A. Bertero. Rna biogenesis instructs functional inter-chromosomal genome architecture. Frontiers in Genetics, 12:645863, 2021.

A. Bertero, P. A. Fields, V. Ramani, G. Bonora, G. Y. mcı H. Reinecke, L. Pabon, W. S. Noble, J. Shendure, and C. E. Murry. Dynamics of genome reorganization during human cardiogenesis reveal an RBM20dependent splicing factory. Nature Communications, 10(1):1538, 2019.

A. Bhaskara, M. Charikar, E. Chlamtac, U. Feige, and A. Vijayaraghavan. Detecting high log-densities: an O(n^¼^) approximation for densest k-subgraph. In Proceedings of the forty-second ACM symposium on Theory of computing, pages 201–210, 2010.

P. Bhat, D. Honson, and M. Guttman. Nuclear compartmentalization as a mechanism of quantitative control of gene expression. Nature Reviews Molecular Cell Biology, 22(10):653–670, 2021.

C. Bulathsinghalage and L. Liu. Network-based method for regions with statistically frequent interchromosomal interactions at single-cell resolution. BMC Bioinformatics, 21(14):1–15, 2020.

E. M. Bunnik, K. B. Cook, N. Varoquaux, G. Batugedara, J. Prudhomme, A. Cort, L. Shi, C. Andolina, L. S. Ross, D. Brady, D. A. Fidock, F. Nosten, R. Tewari, P. Sinnis, F. Ay, J.-P. Vert, W. S. Noble, and K. G. L. Roch. Changes in genome organization of parasite-specific gene families during the Plasmodium transmission stages. Nature Communications, 15(9):1910, 2018.

M. Charikar. Greedy approximation algorithms for finding dense components in a graph. In International workshop on approximation algorithms for combinatorial optimization, pages 84–95. Springer, 2000.

Y. Chen, Y. Zhang, Y. Wang, L. Zhang, E. K. Brinkman, S. A. Adam, R. Goldman, B. Van Steensel, J. Ma, and A. S. Belmont. Mapping 3d genome organization relative to nuclear compartments using tsa-seq as a cytological ruler. J Cell Biol, 217(11):4025–4048, 2018.

P. R. Cook. A model for all genomes: the role of transcription factories. Journal of molecular biology, 395(1): 1–10, 2010.

T. Cremer and M. Cremer. Chromosome territories. Cold Spring Harb Perspect Biol, 2(3):a003889, 2010.

J. R. Dixon, S. Selvaraj, F. Yue, A. Kim, Y. Li, Y. Shen, M. Hu, J. S. Liu, and B. Ren. Topological domains in mammalian genomes identified by analysis of chromatin interactions. Nature, 485(7398):376–380, 2012.

J. M. Dolezal, A. P. Dash, and E. V. Prochownik. Diagnostic and prognostic implications of ribosomal protein transcript expression patterns in human cancers. BMC Cancer, 18:1–14, 2018.

M. F. Duffy, J. Tang, F. Sumardy, H. H. Nguyen, S. A. Selvarajah, G. A. Josling, K. P. Day, M. Petter, and G. V. Brown. Activation and clustering of a plasmodium falciparum var gene are affected by subtelomeric sequences. The FEBS Journal, 284(2):237–257, 2017.

ENCODE Project Consortium. An integrated encyclopedia of DNA elements in the human genome. Nature, 489:57–74, 2012.

A. M. Fenix, Y. Miyaoka, A. Bertero, S. M. Blue, M. J. Spindler, K. K. Tan, J. A. Perez-Bermejo, A. H. Chan, S. J. Mayerl, T. D. Nguyen, et al. Gain-of-function cardiomyopathic mutations in rbm20 rewire splicing regulation and re-distribute ribonucleoprotein granules within processing bodies. Nature Communications, 12(1):6324, 2021.

M. J. Gardner, N. Hall, E. Fung, O. White, M. Berriman, R. W. Hyman, J. M. Carlton, A. Pain, K. E. Nelson, S. Bowman, I. T. Paulsen, K. James, J. A. Eisen, K. Rutherford, S. L. Salzberg, A. Craig, S. Kyes, M.-S. Chan, V. Nene, S. J. Shallom, B. Suh, J. Peterson, S. Angiuoli, M. Pertea, J. Allen, J. Selengut, D. Haft, M. W. Mather, A. B. Vaidya, D. M. A. Martin, A. H. Fairlamb, M. J. Fraunholz, D. S. Roos, S. A. Ralph, G. I. McFadden, L. M. Cummings, G. M. Subramanian, C. Mungall, J. C. Venter, D. J. Carucci, S. L. Hoffman, C. Newbold, R. W. Davis, C. M. Fraser, and B. Barrell. Genome sequence of the human malaria parasite Plasmodium falciparum. Nature, 419(6906):498–511, 2002.

D. Gibson, R. Kumar, and A. Tomkins. Discovering large dense subgraphs in massive graphs. In Proceedings of the 31st international conference on Very large data bases, pages 721–732, 2005.

H. Gu, H. Harris, M. Olshansky, K. Mohajeri, Y. Eliaz, S. Kim, A. Krishna, A. Kalluchi, M. Jacobs, G. Cauer, et al. Fine-mapping of nuclear compartments using ultra-deep hi-c shows that active promoter and enhancer elements localize in the active a compartment even when adjacent sequences do not. BioRxiv, 2021.

W. Guo, S. Schafer, M. L. Greaser, M. H. Radke, M. Liss, T. Govindarajan, H. Maatz, H. Schulz, S. Li, A. M. Parrish, et al. Rbm20, a gene for hereditary cardiomyopathy, regulates titin splicing. Nature Medicine, 18 (5):766–773, 2012.

A. Hafner and A. Boettiger. The spatial organization of transcriptional control. Nature Reviews Genetics, 24 (1):53–68, 2023.

C. Hoencamp, O. Dudchenko, A. M. Elbatsh, S. Brahmachari, J. A. Raaijmakers, T. van Schaik, Á. Sedenõ Cacciatore, V. G. Contessoto, R. G. van Heesbeen, B. van den Broek, et al. 3d genomics across the tree of life reveals condensin ii as a determinant of architecture type. Science, 372(6545):984–989, 2021.

I. L. Ibarra, N. M. Hollmann, B. Klaus, S. Augsten, B. Velten, J. Hennig, and J. B. Zaugg. Mechanistic insights into transcription factor cooperativity and its impact on protein-phenotype interactions. Nature Communications, 11(1):124, 2020.

M. Imakaev, G. Fudenberg, R. P. McCord, N. Naumova, A. Goloborodko, B. R. Lajoie, J. Dekker, and L. A. Mirny. Iterative correction of Hi-C data reveals hallmarks of chromosome organization. Nature Methods, 9:999–1003, 2012.

J. Joo, S. Cho, S. Hong, S. Min, K. Kim, R. Kumar, J.-M. Choi, Y. Shin, and I. Jung. Probabilistic establishment of speckle-associated inter-chromosomal interactions. Nucleic Acids Research, page gkad 211, 2023.

S. Kaufmann, C. Fuchs, M. Gonik, E. E. Khrameeva, A. A. Mironov, and D. Frishman. Inter-chromosomal contact networks provide insights into mammalian chromatin organization. PloS one, 10(5):e0126125, 2015.

S. Khuller and B. Saha. On finding dense subgraphs. In International colloquium on automata, languages, and programming, pages 597–608. Springer, 2009.

T. Krumm and Z. Duan. Understanding the 3D genome: Emerging impacts on human disease. Seminars in Cell & Developmental Biology, 90:62–77, 2019.

E. Lieberman-Aiden, N. L. van Berkum, L. Williams, M. Imakaev, T. Ragoczy, A. Telling, I. Amit, B. R. Lajoie, P. J. Sabo, M. O. Dorschner, R. Sandstrom, B. Bernstein, M. A. Bender, M. Groudine, A. Gnirke, J. Stamatoyannopoulos, L. A. Mirny, E. S. Lander, and J. Dekker. Comprehensive mapping of long-range interactions reveals folding principles of the human genome. Science, 326(5950):289–293, 2009.

D. Lin, G. Bonora, G. G. Yardımcı, and W. S. Noble. Computational methods for analyzing and modeling genome structure and organization. Wiley Interdisciplinary Reviews: Systems Biology and Medicine, 11(1):e1435, 2019.

S. Lomvardas, G. Barnea, D. J. Pisapia, M. Mendelsohn, J. Kirkland, and R. Axel. Interchromosomal interactions and olfactory receptor choice. Cell, 126(2):403–413, 2006.

E. Markenscoff-Papadimitriou, W. E. Allen, B. M. Colquitt, T. Goh, K. K. Murphy, K. Monahan, C. P. Mosley, N. Ahituv, and S. Lomvardas. Enhancer interaction networks as a means for singular olfactory receptor expression. Cell, 159(3):543–557, 2014.

A. B. McKie, J. C. McHale, T. J. Keen, E. E. Tarttelin, R. Goliath, J. J. van Lith-Verhoeven, J. Greenberg, R. S. Ramesar, C. B. Hoyng, F. P. Cremers, et al. Mutations in the pre-mrna splicing factor gene prpc8 in autosomal dominant retinitis pigmentosa (rp13). Human Molecular Genetics, 10(15):1555–1562, 2001.

A. Mészáros, G. Erdős, and Z. Dosztányi. Iupred2a: context-dependent prediction of protein disorder as a function of redox state and protein binding. Nucleic Acids Research, 46(W1):W329–W337, 2018.

K. Monahan, I. Schieren, J. Cheung, A. Mumbey-Wafula, E. S. Monuki, and S. Lomvardas. Cooperative interactions enable singular olfactory receptor expression in mouse olfactory neurons. Elife, 6:e28620, 2017.

K. Monahan, A. Horta, and S. Lomvardas. Lhx2-and ldb1-mediated trans interactions regulate olfactory receptor choice. Nature, 565(7740):448–453, 2019.

S. Mostafavi, D. Ray, D. Warde-Farley, C. Grouios, and Q. Morris. Genemania: a real-time multiple association network integration algorithm for predicting gene function. Genome Biology, 9:1–15, 2008.

C. S. Osborne, L. Chakalova, K. E. Brown, D. Carter, A. Horton, E. Debrand, B. Goyenechea, J. A. Mitchell, S. Lopes, W. Reik, and P. Fraser. Active genes dynamically colocalize to shared sites of ongoing transcription. Nature Genetics, 36(10):1065–1071, 2004.

C. S. Osborne, L. Chakalova, J. A. Mitchell, A. Horton, A. L. Wood, D. J. Bolland, A. E. Corcoran, and P. Fraser. Myc dynamically and preferentially relocates to a transcription factory occupied by igh. PLoS Biology, 5(8):e192, 2007.

A. Papantonis, T. Kohro, S. Baboo, J. D. Larkin, B. Deng, P. Short, S. Tsutsumi, S. Taylor, Y. Kanki, M. Kobayashi, et al. Tnfα signals through specialized factories where responsive coding and mirna genes are transcribed. The EMBO journal, 31(23):4404–4414, 2012.

S. A. Quinodoz, N. Ollikainen, B. Tabak, A. Palla, J. M. Schmidt, E. Detmar, M. M. Lai, A. A. Shishkin, P. Bhat, Y. Takei, V. Trinh, E. Aznauryan, P. Russell, C. Cheng, M. Jovanovic, A. Chow, L. Cai, P. McDonel, M. Garber, and M. Guttman. Higher-order inter-chromosomal hubs shape 3D genome organization in the nucleus. Cell, 174(3):744–757, 2018.

S. S. P. Rao, M. H. Huntley, N. Durand, C. Neva, E. K. Stamenova, I. D. Bochkov, J. T. Robinson, A. L. Sanborn, I. Machol, A. D. Omer, E. S. Lander, and E. L. Aiden. A 3D map of the human genome at kilobase resolution reveals principles of chromatin looping. Cell, 59(7):1665–1680, 2014.

S. B. Reiff, A. J. Schroeder, K. Kırlı, A. Cosolo, C. Bakker, L. Mercado, S. Lee, A. D. Veit, A. K. Balashov, C. Vitzthum, et al. The 4d nucleome data portal as a resource for searching and visualizing curated nucleomics data. Nature Communications, 13(1):2365, 2022.

N. Servant, N. Varoquaux, B. R. Lajoie, E. Viara, C. J. Chen, J.-P. Vert, E. Heard, J. Dekker, and E. Barillot. HiC-Pro: an optimized and flexible pipeline for Hi-C data processing. Genome Biology, 16:259, 2015.

J. Sreedharan, I. P. Blair, V. B. Tripathi, X. Hu, C. Vance, B. Rogelj, S. Ackerley, J. C. Durnall, K. L. Williams, E. Buratti, et al. Tdp-43 mutations in familial and sporadic amyotrophic lateral sclerosis. Science, 319(5870): 1668–1672, 2008.

I. Tanaka, A. Chakraborty, O. Saulnier, C. Benoit-Pilven, S. Vacher, D. Labiod, E. W. Lam, I. Bieche, O. Delattre, F. Pouzoulet, et al. Zranb2 and syf2-mediated splicing programs converging on ect2 are involved in breast cancer cell resistance to doxorubicin. Nucleic Acids Research, 48(5):2676–2693, 2020.

E. L. Van Nostrand, G. A. Pratt, A. A. Shishkin, C. Gelboin-Burkhart, M. Y. Fang, B. Sundararaman, S. M. Blue, T. B. Nguyen, C. Surka, K. Elkins, et al. Robust transcriptome-wide discovery of rna-binding protein binding sites with enhanced clip (eclip). Nature Methods, 13(6):508–514, 2016.

J. Weston, A. Elisseeff, D. Zhou, C. Leslie, and W. S. Noble. Protein ranking: from local to global structure in the protein similarity network. Proceedings of the National Academy of Sciences, 101(17):6559–63, 2004.

P. E. Wright and H. J. Dyson. Intrinsically disordered proteins in cellular signalling and regulation. Nature reviews Molecular cell biology, 16(1):18–29, 2015.

C.-E. Yu, J. Oshima, Y.-H. Fu, E. M. Wijsman, F. Hisama, R. Alisch, S. Matthews, J. Nakura, T. Miki, S. Ouais, et al. Positional cloning of the werner’s syndrome gene. Science, 272(5259):258–262, 1996.

Y. Zhang, T. Li, S. Preissl, M. L. Amaral, J. D. Grinstein, E. N. Farah, E. Destici, Y. Qiu, R. Hu, A. Y. Lee, et al. Transcriptionally active herv-h retrotransposons demarcate topologically associating domains in human pluripotent stem cells. Nature Genetics, 51(9):1380–1388, 2019.

H. Zheng and W. Xie. The role of 3d genome organization in development and cell differentiation. Nature Reviews Molecular Cell Biology, 20(9):535–550, 2019.

